# Mechanochemical Model of Transcriptional Bursting

**DOI:** 10.1101/802751

**Authors:** A. Klindziuk, B. Meadowcroft, A. B. Kolomeisky

## Abstract

Populations of genetically identical cells generally show a large variability in cell phenotypes, which is typically associated with the stochastic nature of gene expression processes. It is widely believed that a significant source of such randomness is the transcriptional bursting, which is when periods of active production of RNA molecules alternate with periods of RNA degradation. However, the molecular mechanisms of such strong fluctuations remain unclear. Recent studies suggest that DNA supercoiling, which happens during transcription, might be directly related to the bursting behavior. Stimulated by these observations, we developed a stochastic mechano-chemical model of supercoiling-induced transcriptional bursting where the RNA synthesis leads to the buildup of torsion in DNA. This slows down the RNA production until binding of an enzyme gyrase to DNA, which releases the stress and allows for the RNA synthesis to restart with the original rate. Using a thermodynamically consistent coupling between mechanical and chemical processes, dynamic properties of transcription are explicitly evaluated. In addition, a first-passage method to evaluate the dynamics of transcription is developed. Theoretical analysis shows that the transcriptional bursting is observed when both the supercoiling and the mechanical stress-release due to gyrase are present in the system. It is also found that the overall RNA production rate is not constant and depends on the number of previously synthesized RNA molecules. A comparison with experimental data on bacteria allows us to evaluate the energetic cost of supercoiling during transcription. It is argued that the relatively weak mechanochemical coupling allows transcription to be regulated most effectively.

**SIGNIFICANCE:** Transcriptional bursting has been cited as one of the probable causes of phenotypic differences in cells with identical genomes. However, the microscopic origin of noisy dynamics in RNA production remains unclear. We developed a thermodynamically-consistent mechano-chemical stochastic model, which, via explicit calculations of dynamic properties, provides a consistent physical-chemical description of how the supercoiling of DNA together with enzymatic activity of gyrases produce transcriptional bursting. It also allows us to explain that the coupling between mechanical and chemical processes might be the reason for efficient regulation of transcription.

## INTRODUCTION

Transcription is the first step in a complex process of gene expression, which is critically important for the functioning of all living systems. It involves copying the genetic code contained in DNA into complementary messenger RNA molecules. This allows the genetic information to be correctly transferred to every cellular process, and it ensures proper development and regulation of organisms (1, 2). A complex network of biochemical and biophysical processes governs and regulates transcription, and even though many aspects of transcription have been uncovered, some of its molecular mechanisms remain poorly understood (1–4). For example, the role of transcription in gene expression variability is still highly debated (3, 4). It is known that genetically identical cells exhibit a wide spectrum of biochemical and biophysical properties, and it is observed in all biological systems, ranging from bacteria to multi-cellular organisms (1, 3). Such variability is a consequence of underlying stochastic processes in gene regulation. It was suggested then that transcriptional bursting, a phenomenon in which the active periods of the synthesis of RNA molecules alternate with periods of transcriptional silence, might be the dominating source of the observed variability in gene expression (3–7).

To explain large fluctuations during the RNA production, the transcriptional bursting is commonly modeled as a multi-state stochastic process (4, 5, 8–14). In this scenario, the RNA synthesis and degradation rates vary in different stochastic states. This randomness is believed to be the source of the bursting phenomena, and it accounts for recent experimental studies that find a spectrum of transcriptional activity levels in certain genes (12, 15, 16). However, the main unresolved fundamental problem here is to understand the molecular mechanisms that lead to multi-state dynamic behavior observed in the transcriptional bursting.

Several theoretical ideas have been proposed to uncover the microscopic origin of transcriptional bursting (17–24). One of them suggests that the transcriptional bursting is a result of the collective dynamics of multiple RNA polymerases (RNAP) that move along the DNA chain and catalyze the synthesis of messenger RNA molecules (17). It was argued that interactions between RNAP molecules can produce bursting behavior, although the molecular details of the process were not explained. But the currently dominating opinion in this field is that the transcriptional bursting is a consequence of coupling between chemical and mechanical processes (18–24). During transcription, the RNAP moves rotationally following the DNA helical chain, unwinding the double-stranded DNA segment in front and reannealing it behind. In living cells, the DNA molecules are frequently topologically constrained, and this leads to a buildup of positive DNA supercoiling in front of the enzyme molecule and the negative DNA supercoiling behind the enzyme molecule. The negative supercoiling can be quickly released by the enzyme topoisomerase I, which is available in significant concentrations in cells (18). But the removal of positive supercoiling requires the enzyme gyrase, which is present in cells in limited quantities. It was argued that the interplay between the mechanical stress buildup and mechanical stress release via the enzymatic action of gyrases is the source of transcriptional bursting. This led to the development of several theoretical models that were able to clarify some aspects of transcriptional bursting (19–24). However, the current theoretical models provide very simplified phenomenological descriptions of the coupling between chemical and mechanical processes. Besides, an unrealistic assumption of a cutoff in RNA production has been implemented in these approaches. Furthermore, existing theoretical methods cannot quantify the coupling between chemical and mechanical forces that govern the transcriptional bursting phenomena.

In this paper, we investigate a discrete-state stochastic mechano-chemical model of transcriptional bursting where the coupling between chemical and mechanical processes is taken into account using a thermodynamically-consistent approach and no assumption of the cutoff in the RNA synthesis is invoked. Also, we develop a first-passage analysis of transcription, which allows us to evaluate the relevant dynamic and energetic properties of the system. Our explicit calculations, supported by Monte Carlo computer simulations, show that the transcriptional bursting is a result of competition between the mechanical forces due to the stress increase in DNA after the synthesis of RNA and the chemical forces due to the action of gyrases that release the mechanical stress. The multi-state dynamic description arises because of the existence of different levels of mechanical stress during transcription with the supercoiling buildup.

## METHODS

### Theoretical Model

To investigate the molecular mechanisms of the transcriptional bursting, we propose a discrete-state stochastic model as illustrated in Fig. 1. The RNAP molecule starts transcription after associating to the DNA chain. If the gyrase molecule is also bound to DNA, then the mechanical stress can not build up, and fast transcription rates are expected: see Fig. 1a. It is assumed that at these conditions the RNA production rate is *α*, while the RNA degradation rate is *β*. This state of the system is labeled as *ON* (Fig 1). After the gyrase dissociates from DNA with a rate *k*_*off*_, mechanical stress increases proportionally to the number of produced RNA molecules, and this slows down the RNA synthesis rate to *α*/*y* ^*j*+1^. Here *j* is the number of RNA molecules produced after the last unbinding of the gyrase from DNA and *y* = exp(*ϵ*/*k*_*B*_*T*) with E being the energetic cost of supercoiling in an RNA synthesis reaction. The state of the system with production rate *a/y* ^*j*+1^ is labeled as *j* (see Fig 1). The parameter *j* can also be viewed as a measure of the degree of supercoiling on the DNA chain: the larger the *j*, the stronger is the mechanical stress on the system and slower is the RNA synthesis reaction rate. The gyrase can bind back to the DNA chain with a rate of *k*_*on*_, and we assume that this immediately removes all mechanical stress, leading to the normal RNA production. A corresponding kinetic scheme for the discrete-state stochastic model of the transcription bursting is presented in Fig. 1b.

**Figure 1:**
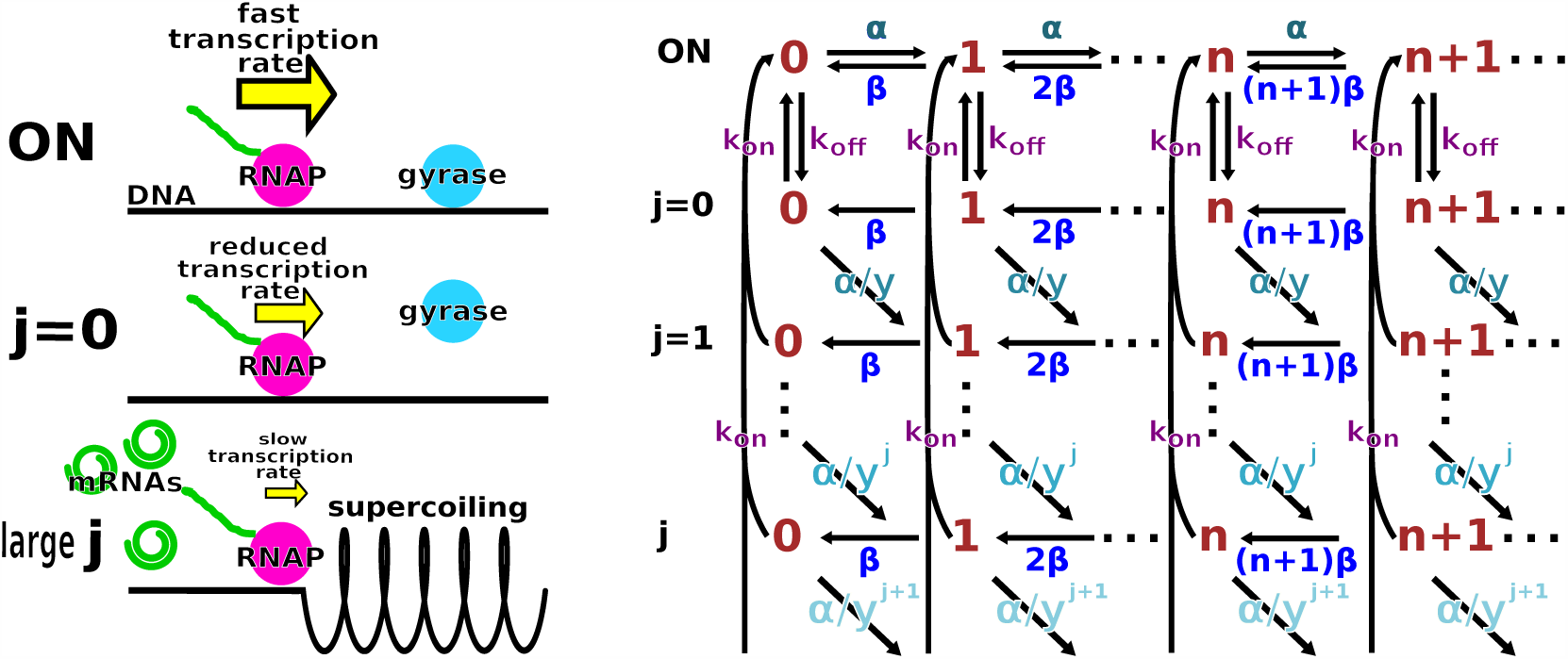
(a) A pictorial depiction of possible kinetic states during transcription. (b) A discrete-state stochastic scheme for the mechano-chemical model of the transcriptional bursting.

It is important to note that our theoretical approach describes the coupling between a chemical (RNA synthesis) processes and a mechanical (supercoiling buildup) process in a thermodynamically consistent way. The energetic parameter *ϵ* describes the additional work that the RNAP molecule must perform when it catalyzes the formation of the messenger RNA molecule in the presence of mechanical stress on DNA. Then following the standard Kramer’s description of chemical rates, this leads to the exponential dependence on the supercoiling energy, *α*/*y*^*j*+1^. Also, in our model, any number of RNA molecules can be produced but the probability of such events exponentially decreases with the degree of mechanical stress in the system. This eliminates the need for introducing non-physical cutoffs.

We define *P*_*ON*_ (*n,t*) as a probability density function to find the system in the state *ON* with *n* produced RNA molecules at the time *t*, and *P*_*j*_ (*n,t*) as a probability density function to find the system in the state *j* (*j* = 0, 1, …) with *n* produced RNA molecules at the time *t*. Then the dynamics of transcription in our model can be described by a set of forward master equations

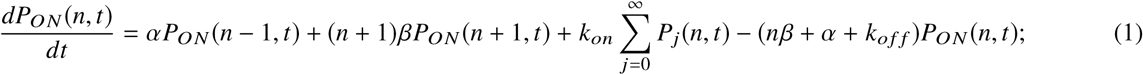

for *j* = 0 we have

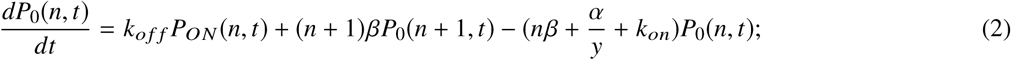

and for *j* > 0

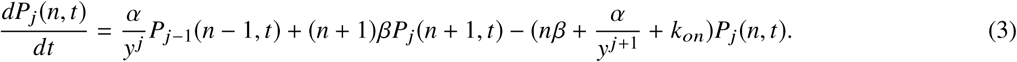

Assuming that the system quickly reaches the steady state (*t* → ∞), dynamical properties of transcription can be evaluated using the method of generating functions (8, 12), as explained in detail in the Supporting Information. It is shown there that the stationary probabilities of kinetic states with different degrees of mechanical stress are given by

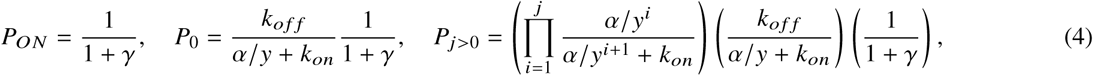

where *γ* = *k*_*off*_*/k*_*on*_ is the gyrase dissociation equilibrium constant. Theoretical calculations also yield the average number of produced RNA molecules (see the Supporting Information),

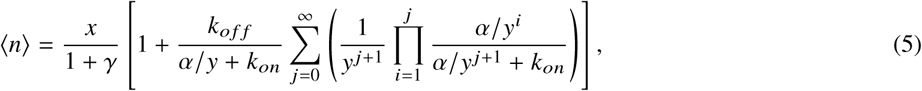

where *x* = *α*/*β* is the equilibrium constant for production/degradation of RNA molecules. In addition, our theoretical method is able to compute the higher moments of the distribution of produced RNA molecules. Specifically, we obtain (see the Supporting Information)

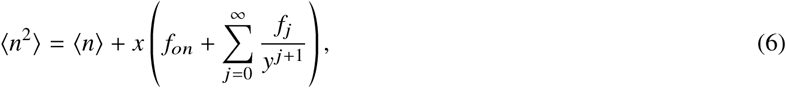

where the coefficients *f*_*ON*_ and *f*_*j*_ are given by

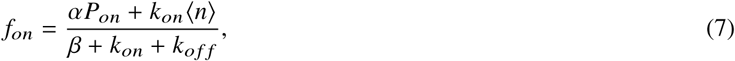

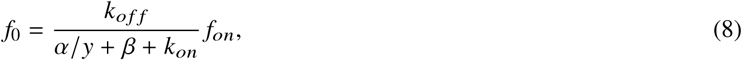

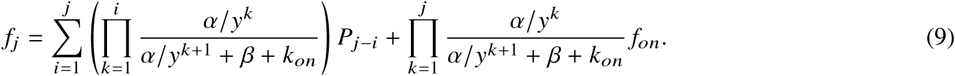

This allows us to evaluate an important parameter, known as a Fano factor,

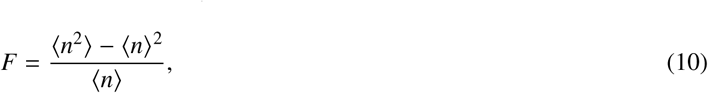

which is a dimensionless measure of the width of the distribution of the produced RNA molecules. It also measures the degree of stochastic noise in the system (5, 10, 12, 18). The Fano factor quantifies the extent of transcriptional bursting in the system. The deviation of this parameter from 1 gives the measure of the burstiness in transcription.

To understand better the complex processes taking place during transcription, we developed a first-passage analysis of the transcriptional bursting, as explained in the detail in the Supporting Information. One can define a new function, 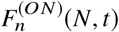, which the probability density function of synthesising exactly *N* RNA molecules at time *t*, given that the system started at *t* = 0 in the state *ON* with *n* RNA molecules already produced. Similarly, we define the first-passage probability function 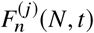 to start from the state *j*. For convenience, we will omit the label *N* in the notations. The temporal evolution of these probability function is governed by the backward master equations,

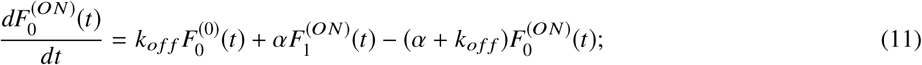

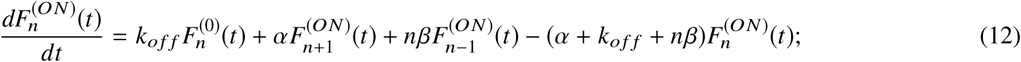

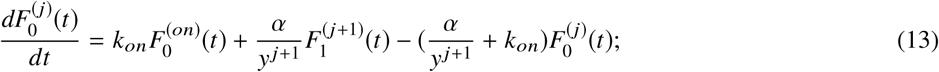

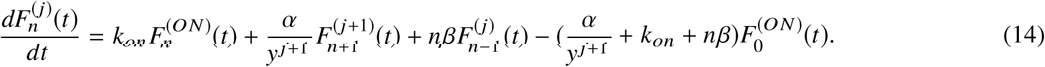

These equations can be analyzed to obtain distributions of production times for *N* RNA molecules, as discussed in the Supporting Information. We are especially interested in mean first-passage times (MFPT), *T*_0_, which are defined as the times to synthesize *N* RNA molecules for the first time starting from the situation without any RNA molecules. These quantities are important because they can be measured in experiments and they also provide important information on the molecular mechanisms of transcription. Explicit analytical results for *T*_0_ can be obtained in several cases, as described in the Supporting Information. We explicitly calculated MFPT in the following situations: 1) when the mechanical stress does not affect the RNA production; 2) when there is supercoiling build up but without gyrases; 3) with supercoiling and with gyrases for *N* = 1; and 4) for general *N* when the gyrase binding/unbinding kinetic rates are much faster than other rates in the system.

In the case where RNA production is unhindered by supercoiling, i.e. when *k*_*off*_ = 0, the system can be viewed effectively as in a single biochemical state (*ON* state). In this case, the MFPT is given by

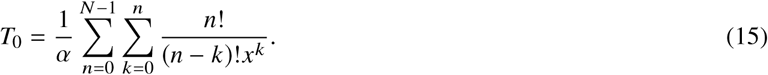

If there is no gyrase present, the supercoiling increases with each new RNA, but the mechanical stress cannot be released. Thus, transcription will eventually stop and it will be unable to resume, and the stationary-state cannot be reached. This corresponds to the mechanochemical model with *k*_*on*_ = 0, and the corresponding MFPT is

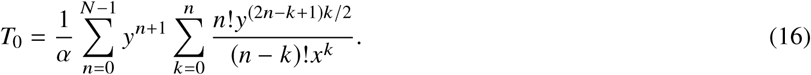

In a general situation, when both mechanical stress and the enzymatic action are present, it can be shown that for *N* = 1

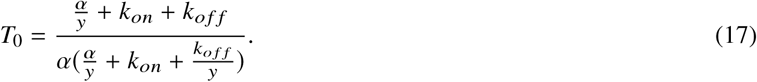

For an arbitrary number of produced RNA molecules *N*, we can calculate analytically MFPT in the limit of fast binding/unbinding rates. Under this condition, the equilibrium is quickly established between different chemical states, leading to

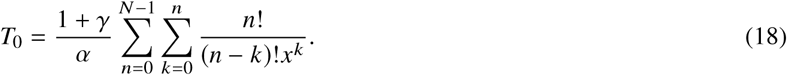

In addition to analytical calculations of the dynamic properties in the discrete-state stochastic model of the transcription bursting, we performed extensive Monte Carlo computer simulations to illustrate better the molecular processes that are taking place in the system. For our calculations we utilized parameters that are similar to those that characterize the transcriptional bursting in bacterial systems (18).

## RESULTS AND DISCUSSION

### Coupling between Mechanical and Chemical Processes in Transcription

The discrete-state stochastic mechanochemical model (Fig. 1b) provides a convenient way to describe the dynamics of transcription and to evaluate the importance of chemical and mechanical processes. Fig. 2 presents the stationary-state distributions and averaged number of produced RNA molecules for different levels of supercoiling affecting the synthesis process. When there is no coupling or the mechanical stress is not created during the RNA production (*ϵ* = 0 and *y* = 1), the distribution is a result of the balance between synthesis and degradation processes. A single maximum is observed in the distribution because in this case the system is found in only one distinct biochemical state, which we labeled as the *ON* state in Fig. 1 (12). But when the mechanical stress builds up influencing the synthesis process, the dynamics in the system changes significantly: see Fig. 2a. The distribution becomes more narrow, and the second peak shifts to the smaller values of *n*. Eventually, for larger supercoiling energetic cost of the synthesis (large *y*) the distribution becomes bimodal, and one of the peaks is at *n* = 0 and another one is at *n* > 0 (Fig. 2a). This exactly agrees with recent experimental observations (18). These results are expected since the mechanical stress on DNA will slow down the synthesis of RNA molecules while the degradation rate will not change. Then the effective production rate, averaged over *ON* and all *j* states will be lower than the *α* rate (for the stress-free situation), and this describes the peak in the distribution for *n* > 0. The peak at *n* = 0 corresponds to those high-stress states (large *j*) in which the RNA synthesis is effectively stopped (*α*/*y*^*j*+1^ → 0). Similar results are observed for the dependence of mean number of produced RNA molecules as a function of the energetic cost of supercoiling: see Fig. 2b. Increasing the coupling between chemical and mechanical processes lowers the output of RNA molecules.

**Figure 2:**
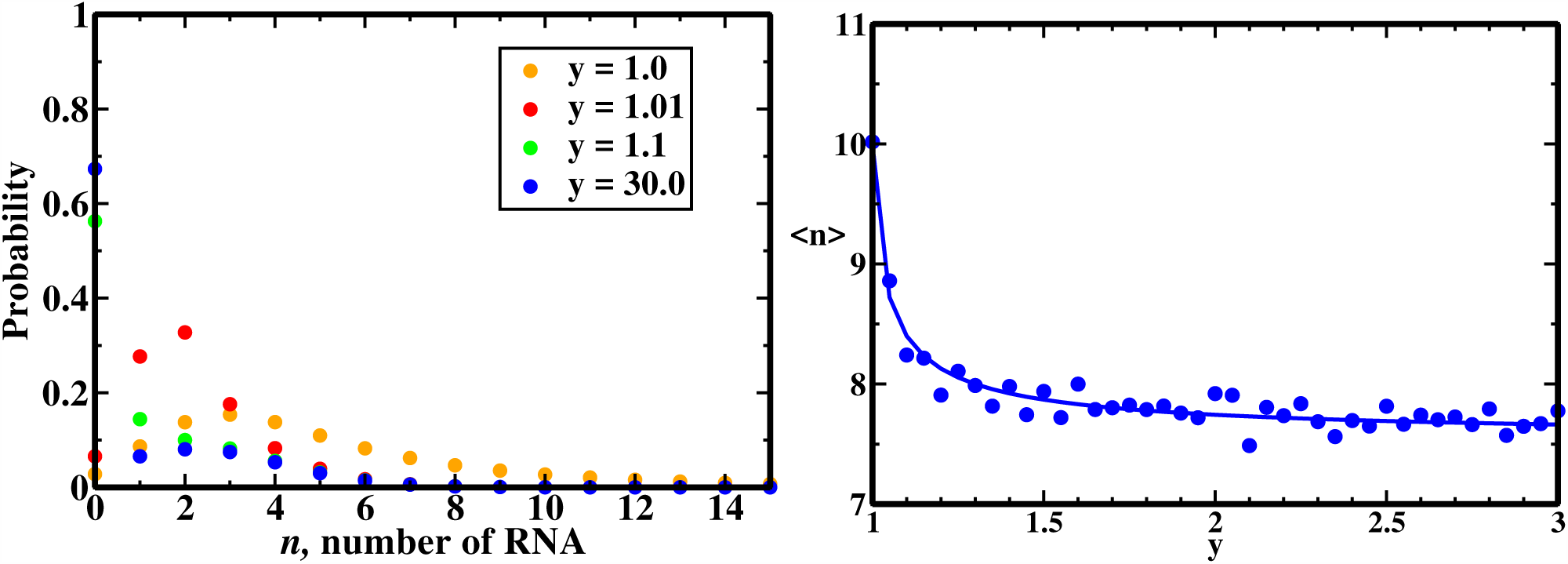
a) Stationary-state distributions for the production of RNA molecules for different energetic costs of supercoiling on synthesis of RNA. The following parameters were used in computer simulations: *α* = 4.0 × 10^−3^ s^−1^, *β* = 1.4 × 10^−3^ s^−1^, *k*_*on*_ = 1.0 × 10^−5^ s^−1^ and *k*_*off*_ = 2.0 × 10^−5^ s^−1^. b) Mean number of produced RNA as a function of the energetic cost of supercoiling. Symbols are from computer simulations and solid line is the analytical result. The following parameters were used in calculations: *α* = 1.0 × 10^−3^ s^−1^, *β* = 1.0 × 10^−4^ s^−1^, *k*_*on*_ = 3.0 × 10^−5^ s^−1^ and *k*_*off*_ = 1.0 × 10^−5^ s^−1^.

Our theoretical method allows us to quantify the degree of supercoiling and how it influences the dynamics in the system. These results are illustrated in Fig. 3 where the stationary-state probability distributions for different states *j* (which are viewed as a measure of mechanical stress on DNA) are presented. In all situations, these distributions are bimodal, reflecting the fact that in the system there is a significant fraction of states with some degree of mechanical stress (*j* > 0) and another important state is the state without supercoiling (*ON* state and *j* = 0). This is a result of a balance between increasing the supercoiling due to the production of more RNA and the slowing down of the synthesis rate for larger *j*. In addition, the binding of the gyrase to DNA leads to the full clearance of mechanical stress in the system. One can see that increasing the gyrase association rate increases the fraction of *j* = 0 state and it shifts the other peak to smaller values of *j* because the system does not have enough time to build up a significant mechanical stress: see Fig. 3a. Varying the energetic cost of supercoiling on the RNA synthesis rate also has a strong effect on the transcription dynamics (Fig. 3b). For weak energetic cost (*y* ∼ 1), the system can easily reach any *j* state because the RNA production is barely affected by the mechanical stress, and this leads to a wide distribution for *j* > 0. For large energetic cost (*y* ≫ 1), only few *j* states can be reached because the synthesis rate decreases very quickly with *j*, and this produces a narrow distribution for small *j*. For intermediate energetic costs, the dynamic behavior that extrapolates between two limiting scenarios is observed (Fig. 3b).

**Figure 3:**
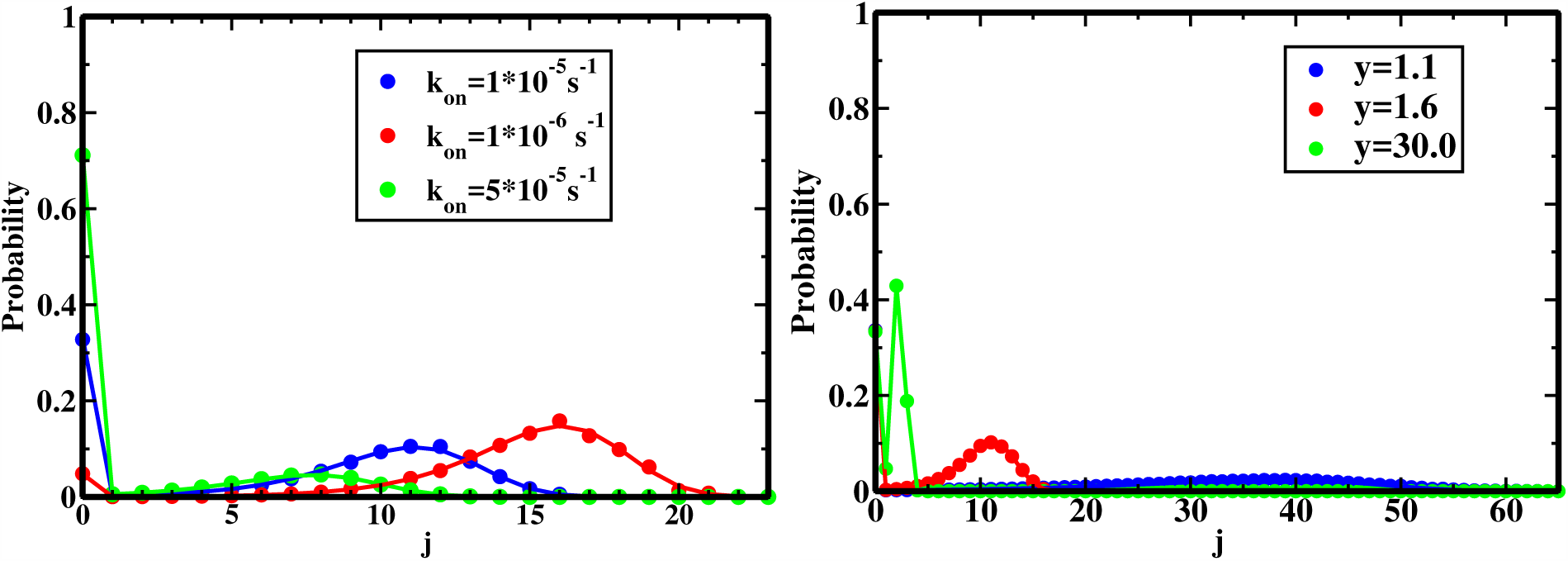
Stationary state probability distributions of mechanical stress on DNA. Symbols are from computer simulations and solid lines are analytical predictions. a) Effect of varying the association rates of gyrases. Parameters used for calculations: *α* = 4.0 × 10^−3^ s^−1^, *β* = 1.4 × 10^−3^ s^−1^” *k*_*off*_ = 2.0 × 10^−5^ s^−1^, and *y* = 1.6. b) Effect of varying the energetic cost of supercoiling. Parameters used for calculations: *α* = 4.0 × 10^−3^ s^−1^, *β* = 1.4 × 10^−3^ s^−1^, *k*_*on*_ = 1.0 × 10^−5^ s^−1^ and *k*_*off*_ = 2.0 × 10^−5^ s^−1^.

An important characteristic of the system is the Fano factor, which provides a dimensionless measure of the bursting behavior (5, 10, 12). It can be calculated explicitly in our theoretical approach, and the results are presented in Fig. 4. The Fano factor *F* correlates with the mean number of produced RNA molecules, ⟨*n*⟩. The mechanical stress lowers the Fano factor, which is expected since the supercoiling slows down the RNA synthesis rate and decreases the number of possible *j* > 0 states that the system can visit. However, the effect is relatively minor (Fig. 4a). A much stronger effect is predicted for varying the gyrase DNA-binding kinetic rates. The larger the association rate *k*_*on*_, the smaller the degree of stochastic fluctuations in the system (the Fano factor is approaching one) because the system spends most of the time in a stress-free *ON* state.

**Figure 4:**
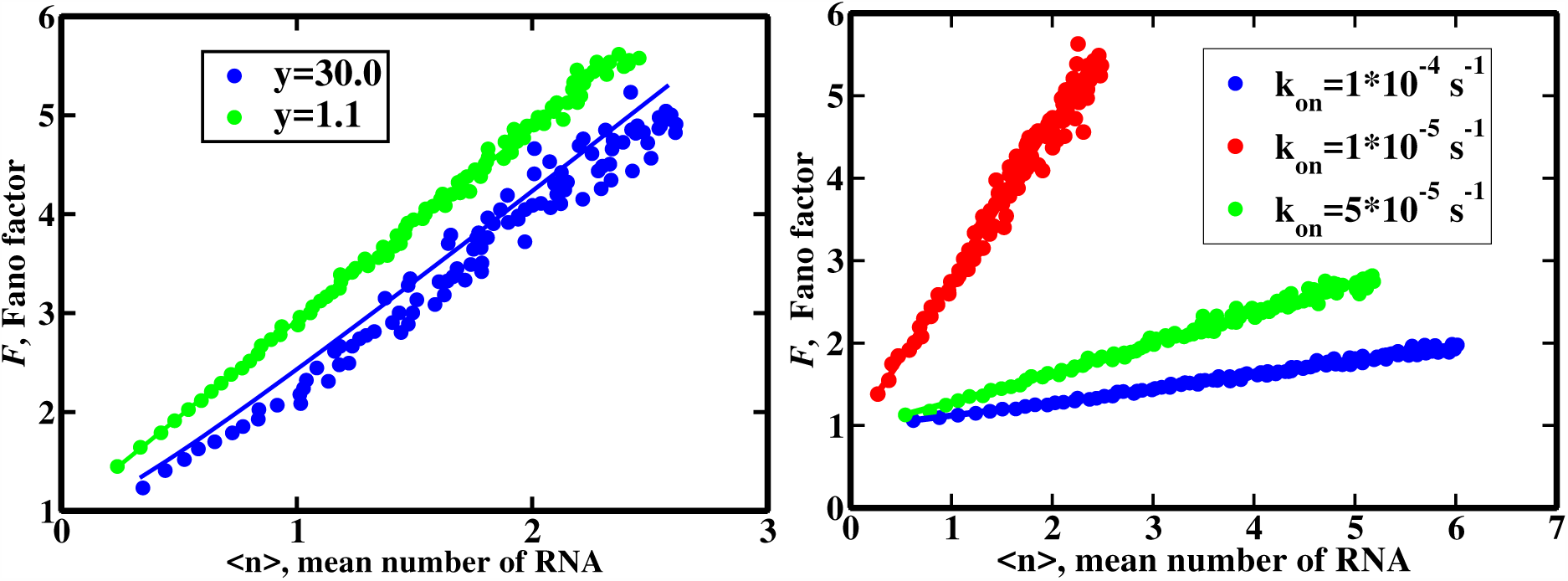
Correlations between the Fano factor and the mean number of produced RNA molecules. Symbols are from computer simulations and solid lines are analytical predictions. a) Effect of varying the energetic cost of supercoiling. Parameters used for calculations: 1 × 10^−3^ < *α* < 1 × 10^−2^ s^−1^, *β* = 1.4 × 10^−3^ s^−1^, *k*_*on*_ = 1 × 10^−5^ s^−1^ and *k*_*off*_ = 2 × 10^−5^ s^−1^. (b) Effect of varying the gyrase association rate to DNA. Parameters used for calculations: 1 × 10^−3^ < *α* < 1 × 10^−2^ s^−1^, *β* = 1.4 × 10^−3^ s^−1^” *k*_*off*_ = 2 × 10^−5^ s^−1^ and *y* = 1.6.

Theoretical analysis of transcription using the discrete-state stochastic model suggests that the bursting behavior is a result of complex interplay between chemical and mechanical forces. The bursting requires having several distinct chemical states with different synthesis and degradation rates. The mechanical stress influences the RNA synthesis rate, and this effect correlates with the number of already produced RNA molecules. This leads to the existence of multiple chemical kinetic states in the system. But to maintain the stationary dynamic behavior, the effect of mechanical stress should be occasionally removed, and this is done by the action of gyrase. Theoretical predictions presented in Figs. 3 and 4 fully support these arguments.

### First-Passage Analysis

Many aspects of the transcription bursting can be better understood by applying the first-passage analysis, which is a powerful tool for investigating complex processes in chemistry and biology (25, 26). The advantage of this method is that it can be applied even when the stationary state is not achieved or cannot be reached at all. Here we estimate the mean times to produce specific quantities of RNA molecules and show how these quantities vary with changing the relevant parameters in the system. The importance of these calculations is that MFPT can be measured in single-molecule experiments (18). The results of our theoretical calculations are presented in Figs. 5 and 6.

**Figure 5:**
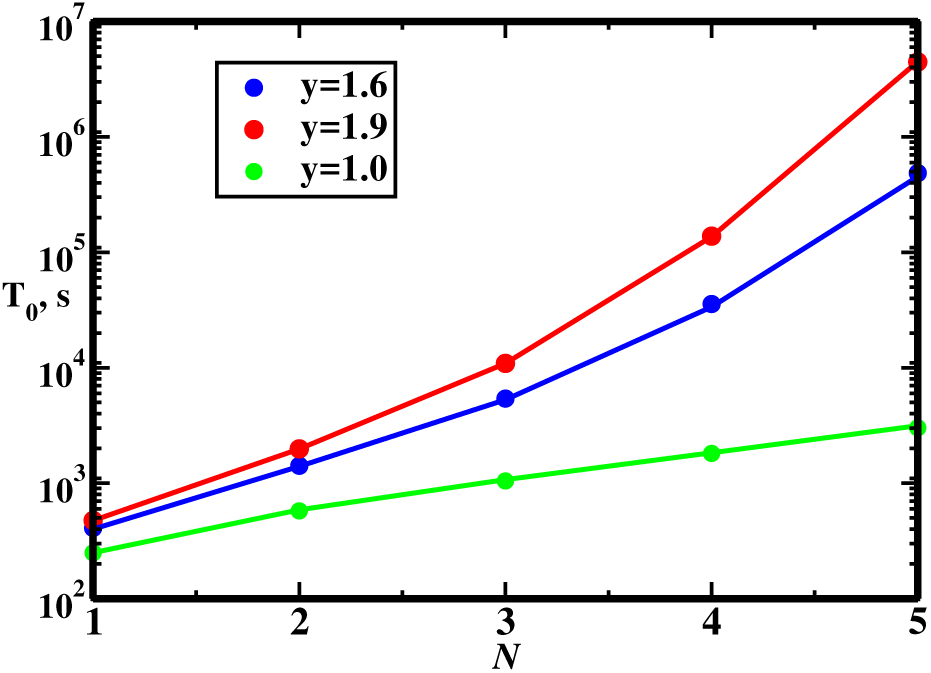
Mean first passage times to produce *N* RNA molecules without mechanical-stress relieving enzymatic action of gyrases for different energetic. costs of supercoiling. Symbols are from computer simulations and solid lines are analytical predictions. Parameters used in calculations: *α* = 4.17 × 10^−3^ s^−1^ and *β* = 1.4 × 10^−3^ s^−1^.

**Figure 6:**
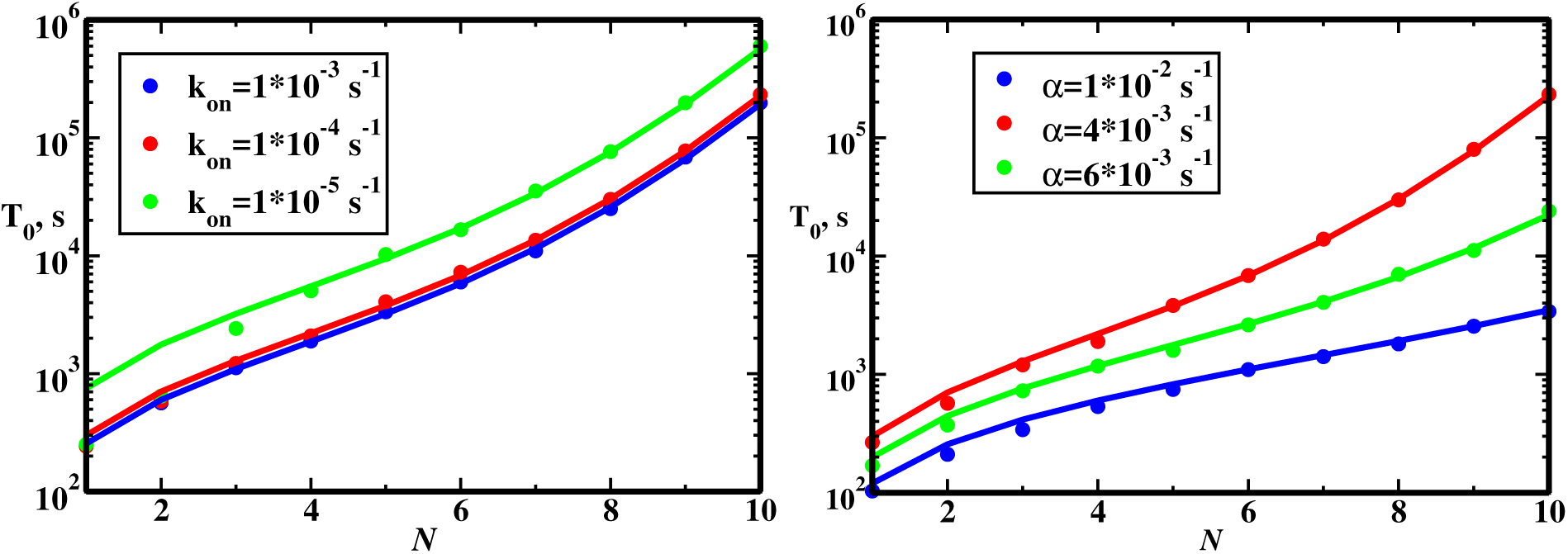
Mean first-passage times to produce *N* RNA molecules in the presence of gyrases. Symbols are from computer simulations and solid lines are analytical predictions. a) The effect of varying the gyrase association rates. Parameters used in calculations: *α* = 4.17 × 10^−3^ s^−1^, *β* = 1.4 × 10^−3^ s^−1^, *y* = 1.6 and *k*_*off*_ = 2.0 × 10^−5^ s^−1^. b) The effect of varying the RNA synthesis rate. Parameters used in calculations: *β* = 1.4 × 10^−3^ s^−1^, *y* = 1.6, *k*_*off*_ = 2.0 × 10^−5^ s^−1^ and *k*_*on*_ = 1.0 × 10^−4^ s^−1^.

We start with the simplest situation when only the supercoiling build-up is taking place, and the gyrase is not present in the system. In this case, it is not possible to remove the mechanical stress and eventually all RNA molecules will be degraded and no stationary transcription bursting will be observed. However, various transient processes are taking place, and the first-passage method can describe them. Fig. 5 shows the MFPT of creating exactly *N* RNA molecules in such a system for different couplings between chemical and mechanical processes. When there is no mechanochemical coupling (*y* = 1), the MFPT grows almost linearly with *N* for synthesis rates faster than the degradation rates. This can be explained by analyzing Eq. (15), which for *x* ≫ 1 (*α* ≫ *β*) predicts (see the Supporting Information) that

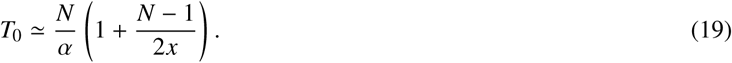

Then, at these conditions, the RNA production rate *R* = *N/T*_0_ is almost constant. However, the situation is different when mechanical stress is influencing the synthesis rate (*y* > 1). Increasing the energetic cost of supercoiling makes the system spend more time producing the same number of RNA molecules. In addition, the MFPT now depends very non-linearly on the parameter *N*. This can be seen in the limiting case of *x* ≫ 1 when Eq. (15) yields,

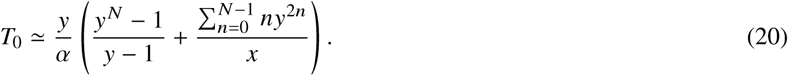

For this case, the RNA production is not a constant and it is sensitive to the degree of mechanical stress in the system.

Let us consider now a more general situation when both mechanical and chemical forces are at play and the transcription bursting can be observed in the stationary state. In this case, the slowing effect of supercoiling is compensated by the enzymatic action of gyrase that removes the mechanical stress. This affects the MFPT of producing RNA molecules in a manner shown in Fig. 6. Increasing the gyrase association rates lowers the mean production times, but the effect is relatively weak for small values of *N* (Fig. 6a). In this case, the gyrase spends more time on DNA, preventing the mechanical stress formation, and this accelerates the overall production of RNA. One should also notice that deviations between theoretical predictions and computer simulations for small values of *k*_*on*_ and for small *N* are due approximate nature of Eq. (20) used in theoretical calculations, where this expression does not work well. Increasing the RNA synthesis rate *α* also lowers the mean production times, but here the effect is stronger for larger *N* values (Fig. 6b). In this case, even if supercoiling is reducing the production rates, the absolute effect is smaller for initially high synthesis rates.

The first-passage analysis can be applied for analyzing recent *in vitro* observations on transcription bursting (18). In these experiments, a single-molecule method has been developed to follow transcription on individual bacterial DNA templates in real time. It was found that in the presence of supercoiling T7 transcription elongation speed (inverse of the MFPT to produce *N* = 1 RNA molecule) slowed down by 38%, while the same measurements for *E*.*coli* produced 47% decrease in the transcription elongation speed (18). In both cases, the gyrase was not present in the system. In our language, *T*_0_(*N* = 1, *y*)/*T*_0_(*N* = 1, *y* = 1) gives the degree of slowing down the transcriptional speed by the mechanical stress. From Eqs. (19) and (20) we obtain

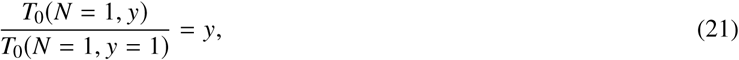

which allows us to evaluate the mechanochemical coupling in these bacteria. It is found that for T7 system, *y* = 1.61 and the energetic cost is *ϵ* = 0.48*k*_*B*_*T*. Similar calculations for *E*.*coli* produce *y* = 1.89 and *ϵ* = 0.64*k*_*B*_*T*. Thus, our analysis suggests that the supercoiling has a relatively weak effect on the transcription of a single RNA molecule. Yet, increasing the mechanical stress by making several RNAs eventually decreases the synthesis rate so much that the RNA degradation starts to dominate. From a biological point of view, this presents a very efficient possible mechanism of transcription bursting regulation via tuning chemical and mechanical processes’ coupling. Very strong coupling would lead to no production of RNA at all, while without coupling no regulation would be possible.

## CONCLUSION

We developed a discrete-state stochastic model of transcription that allowed us to investigate the microscopic origin of transcription bursting phenomena. Our theoretical method takes into account the most relevant features of transcription such as the build-up of supercoiling after each RNA molecule is produced, which slows down the RNA synthesis, and the removal of mechanical stress by gyrase. The model is solved analytically and theoretical calculations are also supported by extensive Monte Carlo simulations. In addition, a novel first-passage analysis of the transcription processes is presented. Our theoretical analysis shows that at the molecular level the transcriptional bursting is a result of competition between chemical processes (RNA synthesis and the gyrase binding to DNA) and the mechanical processes (DNA supercoiling). The presence of the mechanochemical coupling in this system creates several distinct biochemical states with different RNA synthesis and degradation rates. This explains the stochasticity of transcriptional bursting as observed in biological systems. Our first-passage analysis provides the estimate for the degree of mechanochemical coupling in bacterial systems. It is found that the energetic cost of supercoiling on the RNA synthesis rate is relatively weak, but it was argued that this probably leads to very efficient regulation of transcription by tuning the degree of mechanochemical coupling. Thus, the presented theoretical approach clarifies many aspects of complex processes that are taking place during transcription.

Although our theoretical method is able to capture some features of the transcriptional bursting, it is important to discuss its limitations. In our method, the RNA synthesis is viewed as a one-step chemical transition, but in reality, it involves multiple chemical steps with additional assistance from other protein molecules (1, 2). It was assumed also that after the gyrase molecule binds to DNA the mechanical stress is immediately released. However, this probably is not very realistic, and recent experimental measurements (18) and theoretical arguments (23) suggest that the stress relief a relatively slow process. Our theoretical method can be extended to take this effect into account, and we believe that this will not change the main conclusions of our analysis. In addition, in real cells, multiple RNAP molecules simultaneously move on the DNA strand during transcription, but our approach considers only a single-molecule transcription process. It will be important to extend the theoretical analysis in this direction. However, despite these limitations, the presented theoretical results provide a clear molecular picture of transcriptional bursting that may stimulate new experiments and more advanced theoretical studies.

## AUTHOR CONTRIBUTIONS

A.B.K. designed research. B.M. and A.K. and performed research. All authors wrote the article.

## ACKNOWLEDGMENTS

The work was supported by the Welch Foundation (C-1559), the NSF (CHE-1664218), and the Center for Theoretical Biological Physics sponsored by the NSF (PHY-1427654). We also would like to thank Dr. Ido Golding for useful comments and discussions.

